# Chronological lifespan correlates with virulence within *Candida albicans* but long lifespan is not restricted to pathogenic *Candida* species

**DOI:** 10.64898/2026.09.16.751890

**Authors:** Alberto Luna-Carrera, Lilian Rodríguez-Flores, Ericka Moreno-Méndez, Eugenio Mancera, Alexander DeLuna

**Affiliations:** Departamento de Ingeniería Genética, Unidad Irapuato, Centro de Investigación y de Estudios Avanzados del Instituto Politécnico Nacional (Cinvestav), 36824 Irapuato, Mexico; Centro de Investigación sobre el Envejecimiento, Centro de Investigación y de Estudios Avanzados del Instituto Politécnico Nacional (Cinvestav), 07360 Mexico City, Mexico

**Keywords:** Chronological lifespan, Caloric restriction, Fungal virulence, Comparative yeast biology, Clinical isolates, *Candida*

## Abstract

Yeast cells can remain viable for extended periods after entering stationary phase, a survival trait measured as chronological lifespan (CLS). Although CLS is well characterized in *Saccharomyces cerevisiae* as a model of postmitotic aging, its relevance to human-associated fungi remains poorly understood. *Candida* species of the CTG (CUG-Ser1) clade vary widely in ecology and clinical importance, raising the question of whether stationary-phase survival is linked to pathogenicity across species or among strains. Here, we combine an optimized CFU-based CLS assay with scalable flow cytometry of membrane integrity across *Candida* species and genetically diverse *Candida albicans* isolates. CLS varies extensively across species, yet caloric restriction extends survival in every species examined, revealing a remarkably consistent response to reduced glucose despite marked differences in baseline lifespan. Pathogenic species tend toward longer CLS, but long lifespan is not restricted to them, with the non-pathogenic *C. sojae* among the longest-lived species examined. Among 20 genetically diverse *C. albicans* clinical isolates, longer CLS is consistently associated with greater virulence. Our study establishes a methodological framework for comparative CLS analysis, reveals a broadly shared response to caloric restriction, and shows that the association between CLS and virulence emerges within *C. albicans* but not across *Candida* species.

**Highlights:**

- A dual-method framework enables scalable comparative analysis of CLS in *Candida* species and strains.
- CLS varies widely across *Candida*, while caloric restriction extends lifespan in all species examined.
- Long lifespan is not restricted to pathogenic *Candida* species.
- Longer lifespan is associated with higher virulence in *Candida albicans*.

## 1. Introduction

Yeasts frequently encounter conditions in which proliferation is limited but survival remains essential. When nutrients become unavailable, vegetative cells enter stationary phase, a non-dividing state in which long-term viability depends on metabolic remodeling, stress resistance, and the capacity to resume growth when conditions improve (Werner-Washburne et al., 1993). Chronological lifespan (CLS) quantifies this stationary-phase survival by measuring how long cells remain viable over time. In the common yeast, *Saccharomyces cerevisiae*, CLS has been extensively used as a model of postmitotic cellular aging and has revealed conserved links between nutrient sensing, stress responses, and long-term survival (Campos et al., 2018; Cruz-Bonilla et al., 2025; Fabrizio and Longo, 2007). Importantly, CLS varies extensively among yeast genetic backgrounds. Studies of natural *S. cerevisiae* isolates and genetically diverse yeast populations have shown substantial heritable variation in stationary-phase survival, including strain-specific responses to environmental conditions (Barré et al., 2020; De Chiara et al., 2022; Jung et al., 2018).

Less is known about how stationary-phase survival varies in fungi beyond *S. cerevisiae*, including host-associated and opportunistic pathogens. This gap is particularly relevant for *Candida* species, especially those in the CTG (CUG-Ser1) clade, which exhibit remarkable diversity in their ecological niches, stress tolerance, metabolic strategies, and clinical relevance ranging from industrial yeasts to opportunistic human pathogens (Begum et al., 2022; Beyer et al., 2026; Marsaux et al., 2024; Schutz et al., 2024; World Health Organization, 2022). Entry into stationary phase involves extensive physiological remodeling, including metabolic rewiring, activation of stress-response pathways, and increased tolerance to adverse conditions (Cao et al., 2016; den Ridder et al., 2023; Greenlaw et al., 2024; Jacquel et al., 2021; Kusch et al., 2008). In *Candida*, non-proliferative or low-proliferation states have been associated with resistance to oxidative, thermal, and nutrient stresses(Cuéllar-Cruz et al., 2008; Fiori et al., 2012; Richards et al., 2010), traits that may also be relevant during host association, where fungal cells encounter nutrient limitation (Pountain et al., 2021; Rai et al., 2024), immune-mediated pressures(Miramón et al., 2023), and antimicrobial or physicochemical stress (Arastehfar et al., 2023; Lopes et al., 2018; Pierre et al., 2023). Thus, CLS provides a quantitative way to compare stationary-phase survival across *Candida* species and to ask whether stationary phase survival relates to pathogenicity.

Nutrient availability strongly influences the transition from growth to stationary-phase survival. In *S. cerevisiae*, caloric restriction, typically achieved by reducing glucose concentration without inducing starvation, is a well-established intervention extending CLS (Wei et al., 2008). Reduced glucose alters nutrient-sensing pathways including TOR, PKA, Sch9, and the filamentous-growth MAPK pathway, linking nutrient availability to metabolic remodeling, stress resistance, and long-term viability (Aluru et al., 2017; Campos et al., 2018; Karunanithi and Cullen, 2012; Mohammad et al., 2021; Zhang et al., 2025). Yeast species also differ substantially in glucose metabolism, particularly in the balance between respiration and fermentation, and *Candida* species generally retain respiratory activity under glucose-rich conditions that repress respiration in *S. cerevisiae* (Christen and Sauer, 2011; Niimi et al., 1988). In *Candida*, however, reduced glucose has not been systematically examined as a modifier of stationary-phase survival. Testing CLS under caloric restriction therefore asks whether this canonical longevity response extends beyond *S. cerevisiae*.

Across species, any relationship between stationary-phase survival and pathogenicity may be obscured by deeper evolutionary divergence and by species-specific physiology. A complementary question is therefore whether CLS variation is associated with pathogenicity within a single species background. In opportunistic pathogenic fungi, there is considerable variation in virulence among isolates of the same species, including differences in stress tolerance and other traits that influence survival during infection(Day et al., 2018; Shankar et al., 2020). *Candida albicans*, one of the most prevalent human fungal pathogens, exhibits pronounced genetic and phenotypic diversity among clinical isolates, including differences in virulence in experimental infection models (Davari et al., 2026; Freese and Beyhan, 2023; Marcos-Zambrano et al., 2020; Wu et al., 2007). This makes *C. albicans* a suitable system to test whether stationary-phase survival is linked to pathogenicity at the intraspecific scale.

A key challenge in comparative CLS analysis is choosing a viability readout that is both informative and scalable. This issue has been recognized in the CLS field, where different assay formats can emphasize distinct aspects of survival and complicate comparisons across datasets(Cruz-Bonilla et al., 2025; Pereira and Saraiva, 2013; Smith et al., 2016). Among commonly used approaches, classical CLS assays rely on colony-forming units (CFUs), which measure the ability of aged cells to resume proliferation. Although these assays remain the standard readout, they are labor-intensive and can be variable across strain panels. In contrast, flow cytometry-based assays use fluorescent staining to estimate viability from single-cell membrane-permeability profiles offering a higher-throughput complementary readout (Ocampo and Barrientos, 2011; Tang et al., 2025). Whether these approaches provide equivalent CLS estimates in *Candida* species, also across conditions, remains unclear.

In this study, we established a comparative CLS framework by combining optimized CFU-based assays with flow cytometry live/dead measurements to assess stationary-phase survival in *Candida*. We applied this framework across *Candida* species and among genetically diverse *C. albicans* clinical isolates with available virulence estimates in a murine infection model (Wu et al., 2007). Comparative profiling revealed extensive CLS diversity across species, while caloric restriction extended survival in every species examined. Pathogenic species tended toward longer CLS, but long lifespan was not restricted to them. Within *C. albicans*, longer CLS correlated with greater virulence among genetically diverse isolates. Together, these findings identify a broadly shared capacity for CLS extension under caloric restriction and show that, although long CLS is not specific to pathogenic *Candida* species, quantitative variation in CLS correlates with virulence within *C. albicans*.

## 2. Materials and Methods

### 2.1. Yeast strains and media

The multispecies panel (**Table 1**) included *S. cerevisiae*, *C. albicans*, *C. parapsilosis*, *C. dubliniensis*, *C. maltosa*, *C. sojae*, *C. tropicalis*, *Candidozyma auris* (syn. *Candida auris*), and *Nakaseomyces glabratus* (syn. *Candida glabrata*). An additional panel of 20 *C. albicans* strains (**Table 2**) was used for the intraspecific analysis of CLS and virulence (Wu et al., 2007). Strains were maintained as frozen stocks at −80 °C and recovered on YPD agar before experimentation.

**Table 1.**
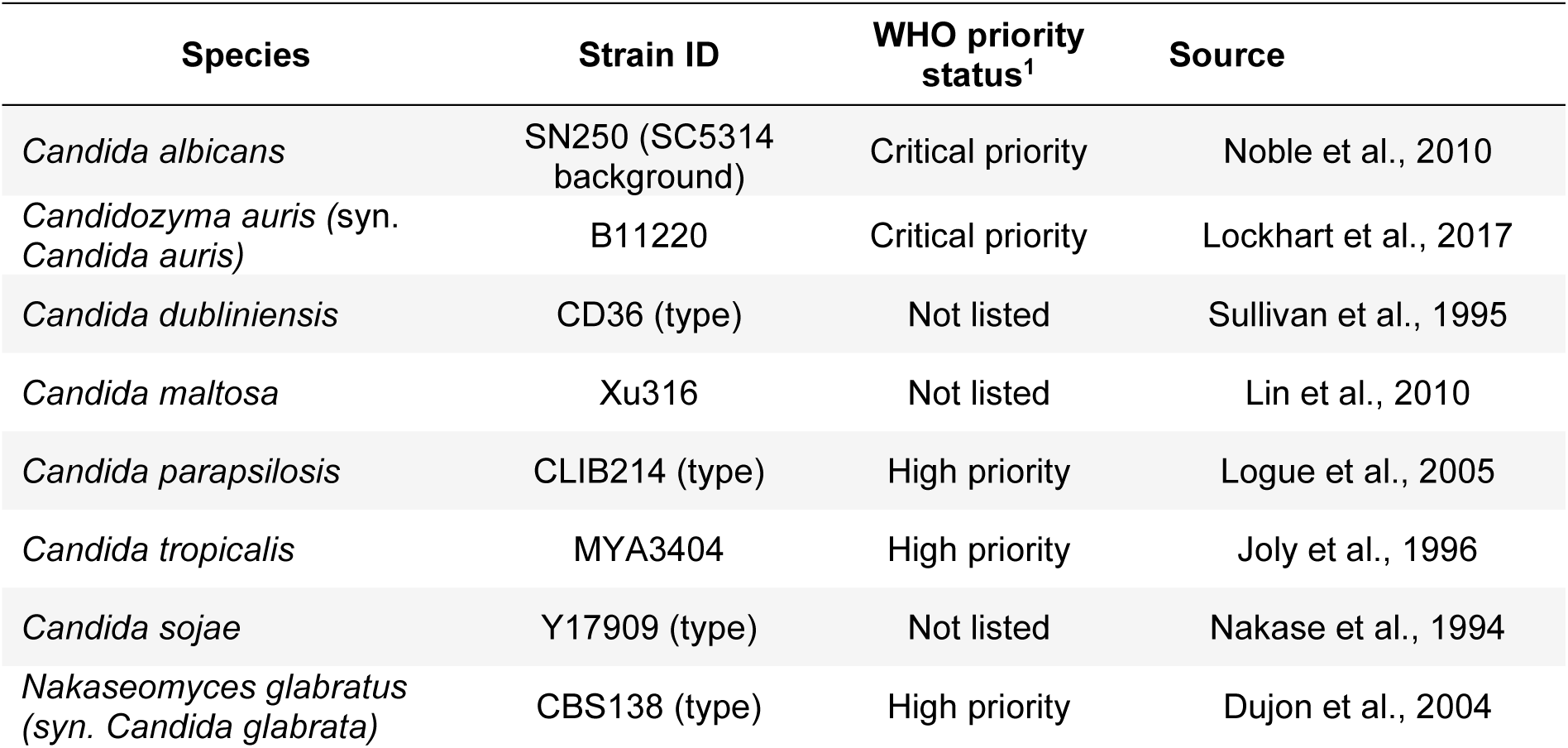
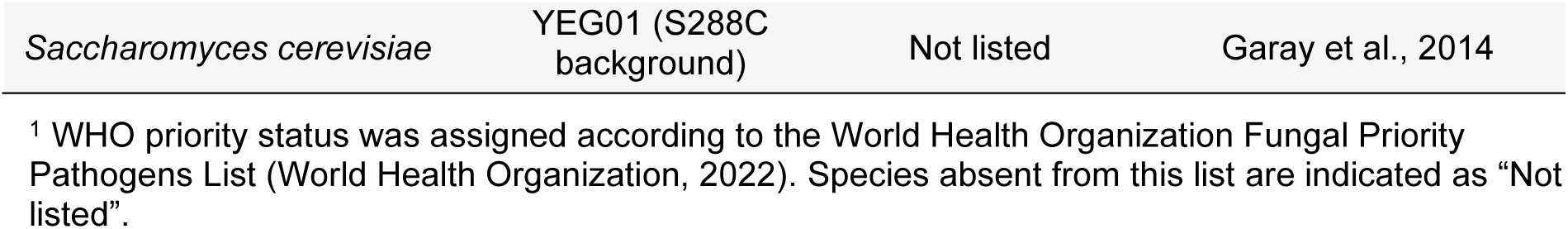
Strains of the yeast species used in this study.

**Table 2.** *Candida albicans* clinical isolates used in this study^1^.

| Strain ID | Virulence Score<sup>2</sup> | Strain ID | Virulence Score<sup>2</sup> |
| --- | --- | --- | --- |
| P75063 | 0.882 | P57072 | 0.382 |
| GC75 | 0.864 | P78042 | 0.373 |
| P37039 | 0.816 | P75016 | 0.313 |
| P76067 | 0.782 | P75010 | 0.296 |
| L26 | 0.736 | P60002 | 0.227 |
| P34048 | 0.650 | P76055 | 0.182 |
| P37005 | 0.525 | 19F | 0.180 |
| P37037 | 0.523 | P87 | 0.059 |
| P57055 | 0.464 | 12C | 0.043 |
| P78048 | 0.45 | P94015 | 0.011 |

<sup>1</sup>All strains are from (Wu et al., 2007).
<sup>2</sup>Virulence scores were calculated as 1 – normalized AUC of the mouse survival curve following inoculation with each *C. albicans* strain (Wu et al., 2007) , such that higher scores correspond to greater virulence.

Growth media used were synthetic complete (SC) and yeast extract–peptone–dextrose (YPD). Unless otherwise stated, percentages are expressed as weight/volume. SC medium contained 0.67% yeast nitrogen base without amino acids, 0.2% amino acid supplement mixture (Yeast Synthetic Drop-out Medium Supplements without uracil, Sigma-Aldrich, Y1501), 0.076 g/L uracil (Sigma-Aldrich, U0750), and either 2% glucose under standard conditions or 0.5% glucose under caloric restriction (Campos et al., 2018). YPD contained 1% yeast extract, 2% peptone, and 2% glucose (Sherman, 2002).

### 2.2. Chronological lifespan culture conditions and sampling

SC was used for CLS assays, with minimal filamentation observed during stationary phase. 30 °C overnight cultures were used to inoculate fresh SC medium at an initial optical density of OD600 = 0.1. These aging cultures were grown in 25 mL of medium in 125-mL Erlenmeyer flasks covered with aluminum foil and incubated at 30 °C with shaking at 200 rpm for up to 60 d. Aging cultures were started under either non-restricted conditions (NR; 2% glucose) or caloric restriction (CR; 0.5% glucose).

For the comparative CLS experiments, aliquots from the same aging cultures were analyzed by CFU assays and flow cytometry. Three independent biological replicates were performed for each strain and condition, each initiated from a distinct culture. Technical replicates were included as specified for each assay. Stationary-phase age 0 was defined as 24 h after inoculation and was used as the baseline for survival measurements. For the comparative species experiments, cultures were sampled daily from days 1 to 8 and then on days 10, 12, 15, 18, 21, 28, 36, 44, 51, and 60 after inoculation. For the intraspecific *C. albicans* analysis, the 20-strain panel was evaluated on days 1, 11, and 16 after inoculation, corresponding to 0, 10, and 15 days of chronological aging, respectively.

### 2.3. Optimized CFU-based CLS determination

Colony forming unit based CLS was determined from the ability of aged cells to form colonies after transferring back to nutrient-rich medium. At each sampling time point, 100 µL of aging culture was serially diluted six times 10-fold in sterile Milli-Q water. For each strain and time point, a dilution yielding between 30 and 300 colonies was selected for plating, as described previously(Pasrija and Kumari, 2025).

CFUs were quantified by spot plating five 40-µL drops of the selected dilution onto YPD agar for each biological replicate. Plates were incubated at 30 °C for 48 h, and colonies were quantified using ImageJ v1.54d (National Institutes of Health, Bethesda, MD, USA). For each biological replicate, CFU counts from five technical replicates were averaged. CFU/mL was calculated by correcting the mean colony count for the plated volume and dilution factor. Survival was normalized to the survival measured at day 0 for each biological replicate, which was set as 100% survival.

### 2.4. Flow cytometry-based CLS determination

Membrane integrity during chronological aging was quantified by flow cytometry using the LIVE/DEAD® FungaLight™ Yeast Viability Kit (Thermo Fisher Scientific) containing SYTO9 and propidium iodide (PI), as described previously(Campos et al., 2018; Meneses-Plascencia et al., 2026). In brief, at each sampling time point, 6 µL of aging culture were mixed with 50 µL of staining solution prepared in BD™ FACSFlow™ buffer by diluting each dye stock 1:1000. The final SYTO9 and PI concentrations were 2.98 µM and 17.9 µM, respectively. Samples were incubated for 15 min at room temperature in the dark with agitation before analysis using a BD LSRFortessa™ flow cytometer equipped with a high-throughput sampler. SYTO9 fluorescence was excited with a 488-nm laser and collected through 505-nm long-pass and 525/50-nm band-pass filters, whereas PI fluorescence was excited with a 561-nm laser and collected through a 586/15-nm band-pass filter. A total of 10,000 events were acquired per technical replicate. Cell populations were identified based on bivariate SYTO9-versus-PI fluorescence plots using species-specific polygon gates, and membrane-intact cells were defined by PI exclusion. Gates were established separately for each species using controls prepared from 24-h cultures: untreated cells as the live-cell control, cells heat-treated at 60°C for 15 min as the dead-cell control, and a 1:1 mixture of untreated and heat-treated cells as the mixed live/dead control. These controls represented nominal 100% live, 50:50 live/dead, and 100% dead reference populations, respectively (Tang et al., 2025). The resulting gates were applied consistently across samples and time points for each species. Data were acquired and analyzed using the FACSDiva software version 8.0.1 (BD Biosciences)

Four technical replicates were analyzed for each biological replicate and averaged before subsequent analyses. Survival was normalized to survival at day 0 for each biological replicate, which was defined as 100% survival. CLS curves were generated from the resulting membrane integrity measurements over time.

### 2.5. CLS data analysis and comparative statistics

For comparative analyses across species, CLS was summarized using the survival integral, calculated as the area under the survival curve (AUC). AUC values were calculated independently for each biological replicate by numerical integration in MATLAB (R2021a) from day 0 to day 60 across all samples. Mean survival integrals were then calculated across biological replicates for each species and condition. Responses to caloric restriction were visualized as reaction norms connecting survival integrals under NR and CR conditions for each species. For the intraspecific *C. albicans* analysis, CLS was reported directly as the percentage of surviving cells at days 10 and 15 of chronological aging, with survival at each time point expressed relative to day 0 for each biological replicate.

### 2.6. Species pathogenicity classification and virulence data extraction and analysis

Species included in the critical- or high-priority groups were designated WHO-listed, whereas species not included in the WHO FPPL were designated not WHO-listed (World Health Organization, 2022). For the purposes of comparative analyses, WHO-listed species were treated as the pathogenic group, using WHO priority status as an operational proxy for pathogenicity. The classification assigned to each species is reported in **Table 1**.

Virulence data for the 20 *C. albicans* strains were obtained from previously published murine systemic infection experiments (Wu et al., 2007). In these experiments, mice were inoculated with individual *C. albicans* strains and host survival was monitored for up to 22 days. Published mouse survival curves were digitized using WebPlotDigitizer v5.2, and the AUC was calculated. AUC values were normalized to the maximum possible survival AUC over the 22-day observation period, yielding values from 0 to 1, with 1 corresponding to survival throughout the observation period. A virulence score was then calculated as:

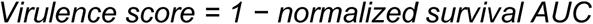

A virulence score of 0 therefore indicates no mortality, whereas higher scores indicate greater mortality and shorter host survival. For strains with independent replicate infection experiments, virulence scores were calculated separately for each replicate survival curve and then averaged to obtain a single strain-level score. The calculated virulence data are provided in **Table 2**.

### 2.7. Statistical analysis

All statistical analyses were performed using MATLAB (R2021a). Comparisons between samples or experimental conditions were performed using two-tailed Student’s *t*-tests using paired tests for matched NR–CR comparisons within the same species or strain and unpaired tests for independent comparisons. Differences in AUC among species under NR were assessed using one-way ANOVA. Technical replicates were averaged within each biological replicate before statistical analysis, with biological replicates used as the statistical unit. Associations between CLS and virulence score across *C. albicans* strains were assessed using Pearson correlation.

## 3. Results

### 3.1. Complementary CFU and flow cytometry assays enable scalable CLS profiling in *Candida*

To examine whether long-term stationary-phase survival is associated with pathogenicity in *Candida*, we developed a comparative CLS framework combining classical CFU-based survival measurements with scalable analyses across species and strains. We established a dual-readout approach in which CFU-based assays and flow cytometry were performed in parallel on the same aging cultures (**Fig. 1a**). This design allowed us to compare colony-forming capacity with a membrane-integrity readout.

**Figure 1.**
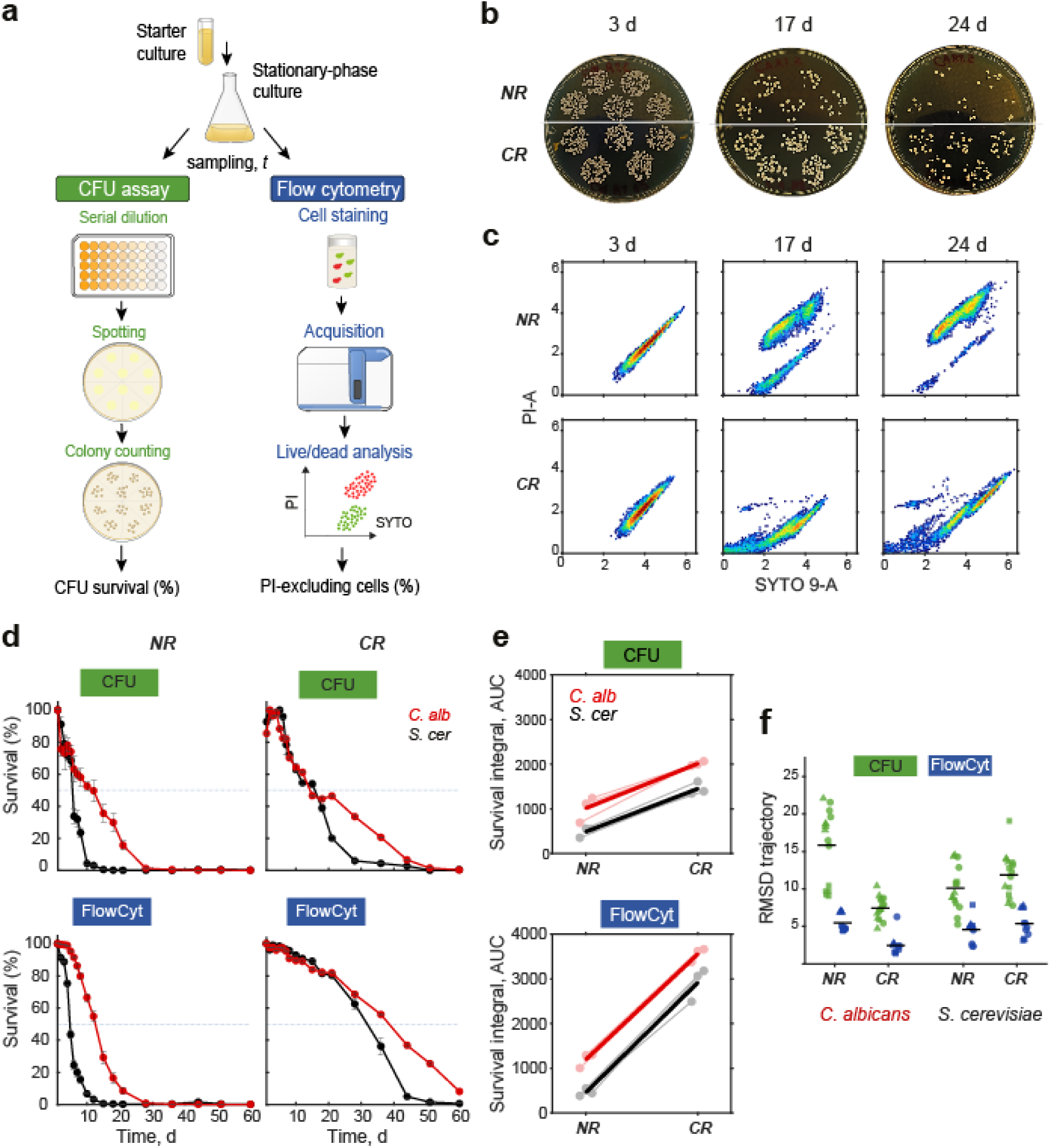
Complementary CFU and flow cytometry assays enable chronological lifespan profiling in *Candida*. **a)** Experimental framework for parallel measurement of CLS by optimized colony-forming unit (CFU) assays and flow cytometry (FlowCyt) based on SYTO9/propidium iodide staining. **b)** Representative CFU plates from *C. albicans* cultures aged under non-restricted (NR, 2% glucose) or caloric-restriction (CR, 0.5% glucose) conditions. **c)** Representative flow-cytometry profiles from the same strain and aging conditions, showing changes in membrane-integrity status over time. **d)** CLS trajectories of *C. albicans* (red) and *S. cerevisiae* (black) measured by CFU and flow cytometry under NR (left) and CR (right). Points represent mean survival across three independent experiments, and error bars indicate the SEM at each time point. **e)** Survival integral under NR and CR for each species and assay. Light lines represent the three independent biological replicates, and solid lines indicate their mean. **f)** Assay variability quantified as the root mean square deviation (RMSD) of individual survival trajectories from the corresponding mean survival curve across sampled time points. Symbols indicate three independent biological replicates (circles, triangles, and squares), each measured in five and four technical replicates for CFU and flow cytometry, respectively; black horizontal lines indicate the mean RMSD.

An important challenge in optimizing CLS assays for *Candida* species was the propensity of some of these yeasts to undergo filamentation. Although both yeast and filamentous morphologies are important for *Candida* colonization and pathogenesis, we focused on conditions that minimized filamentation because filamentous growth can interfere with CFU quantification and complicate flow-cytometric assessment of membrane integrity. We therefore used SC medium to minimize morphological variation as a technical confounder. Under these conditions, *C. albicans* remained predominantly in yeast form, with only minimal filamentation after 16 days in stationary phase (data not shown). Aging cultures were then serially diluted and spot-plated in technical replicate from dilutions yielding countable colonies. In parallel, matched aliquots were stained with SYTO9 and PI and analyzed by flow cytometry to quantify membrane-intact cells based on PI exclusion. Representative time courses showed the expected decline in colony-forming capacity (**Fig. 1b**) and corresponding shifts in flow-cytometry profiles during aging (**Fig. 1c**). Thus, both assays captured the progressive loss of survival over time while measuring distinct biological properties of aged cells.

Using our complementary CLS assays, we asked how chronological survival in *C. albicans* compares with that in *S. cerevisiae* under non-restricted (NR, 2% glucose) and caloric-restriction conditions (CR, 0.5% glucose). Under NR, *C. albicans* survived substantially longer than *S. cerevisiae* in CFU-based measurements, with a half-life of 11.8 days compared with 5.5 days in *S. cerevisiae*; this species difference was also captured by flow cytometry, with half-lives of 4.8 and 12.6 days for *S. cerevisiae* and *C. albicans*, respectively (**Fig. 1d**).

To capture the full survival trajectory for statistical comparisons, we summarized CLS using the AUC. CR extended survival in both species by both readouts, with significant increases in survival AUC. In *S. cerevisiae*, these increases were significant by both CFU and flow cytometry (*p* = 0.004 and *p* = 0.006, respectively), and the same was true for *C. albicans* (*p* = 0.020 and *p* = 3.01 x 10^-5^, respectively; paired two-tailed Student’s *t*-tests).

Notably, absolute survival estimates became method-dependent under CR. In both species, flow cytometry reported higher survival than CFU-based measurements, indicating that PI exclusion and colony-forming capacity become partially decoupled under this condition. Accordingly, the apparent magnitude of CR-mediated lifespan extension was greater when assessed by flow cytometry than by CFU (**Fig. 1e**). Together, these results show that *C. albicans* exhibits a longer NR CLS than *S. cerevisiae*, while CR extends lifespan in both species consistently across assays despite method-dependent differences in absolute survival estimates in this condition.

To assess whether flow cytometry provided sufficient reproducibility for subsequent comparative profiling, we quantified assay variability across independent experiments using the root mean square deviation (RMSD) of each survival trajectory from the corresponding mean curve (**Fig. 1f**). Flow cytometry showed variability on the same general scale as CFU, although reproducibility differed across species and growth conditions. This level of reproducibility, together with the scalability of the assay, supported its use for subsequent comparative CLS profiling. Together, these results support CFU as the classical proliferative-survival reference and flow cytometry as a scalable PI-exclusion readout for relative CLS comparisons across species and strains.

### 3.2. CLS varies extensively across *Candida* species despite a shared response to caloric restriction

With flow cytometry established as a scalable readout for relative CLS comparisons, we applied it to several representatives of the CTG (CUG-Ser1) clade, together with *N. glabratus* (formerly *C. glabrata*) and included *S. cerevisiae* as a comparative reference (**Fig. 2a**). CLS was estimated using flow cytometry under non-restricted conditions (NR, 2% glucose) and caloric restriction (CR, 0.5% glucose). Under NR, species displayed markedly different survival trajectories, ranging from rapid loss of membrane integrity (e.g., *C. dubliniensis*) to prolonged survival over several weeks (e.g., *C. sojae*). The differences between species quantified by the AUC were statistically significant (*p* = 8.83 x 10^-8^; one-way ANOVA) (**Fig. 2b**), indicating substantial diversity in long-term stationary-phase survival across species.

**Figure 2.**
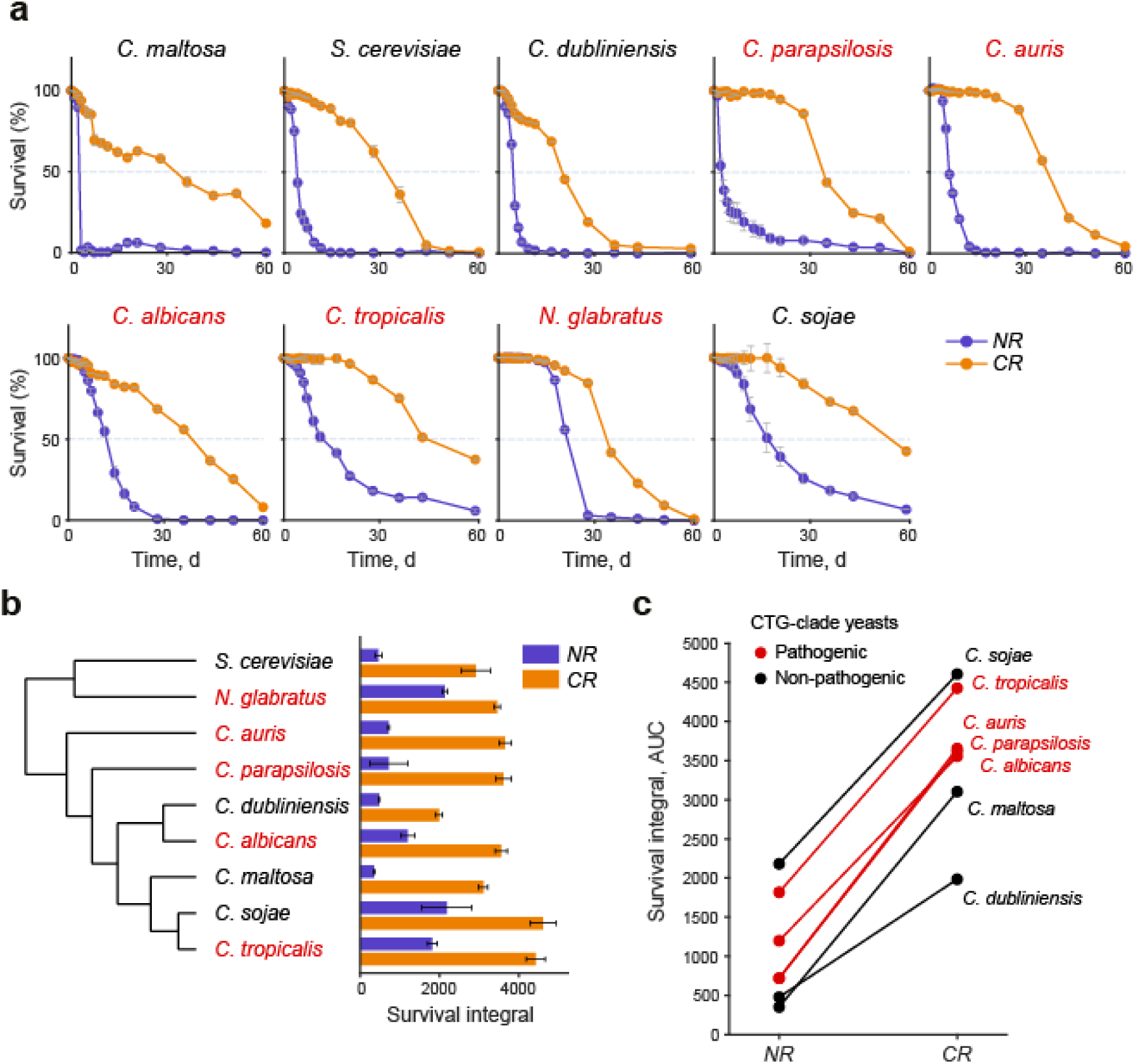
CLS varies extensively across *Candida* species, while caloric restriction consistently extends lifespan. **(a)** Flow cytometry-based CLS profiles for the yeast species analyzed under non-restricted (NR, 2% glucose; purple) and caloric-restriction conditions (CR, 0.5% glucose; orange). Points represent mean survival across three independent experiments, and error bars indicate the SEM at each time point. *S. cerevisiae* was included as a reference and *N. glabratus* as an important yeast pathogen outside the CTG clade. **(b)** Phylogenetic relationships among the species analyzed and their corresponding survival integrals under NR and CR; branch lengths are not proportional to evolutionary distance. Cladogram is based on established phylogenetic relationships among the species analyzed (Chávez-Tinoco et al., 2024; Shen et al., 2016). Bars represent mean survival integrals across three independent experiments, and error bars indicate the SEM. **(c)** Reaction norms showing the change in survival integral from NR to CR for each species. Throughout the figure, pathogenic and non-pathogenic species according to the WHO priority list (Table 1) are shown in red and black, respectively.

Despite the extensive interspecific variation in non-restricted CLS, reduced glucose extended survival in every species examined (**Fig. 2a,b**). Across most species, the magnitude of this response was also remarkably similar, with survival integrals increasing from widely different NR values along approximately parallel reaction norms (**Fig. 2c**). Thus, species largely retained their relative differences in CLS while shifting toward longer survival under CR. One notable exception was *C. dubliniensis*, which combined a short non-restricted CLS with a comparatively weaker response. To verify that these interspecific patterns were also captured by colony-forming capacity, we selected three species spanning distinct CLS and pathogenicity for CFU-based measurements, namely the short-lived non-pathogenic *C. maltosa*, the long-lived non-pathogenic *C. sojae*, and the pathogenic *C. parapsilosis* (**Fig. 3a**). CFU survival curves recapitulated their relative CLS differences under NR, while all three species showed increased survival under CR; reaction-norm analysis similarly reproduced the trends observed by flow cytometry (**Fig 3b**; compare **Fig. 2c**). Overall, these results reveal extensive variation under non-restricted conditions, alongside a shared and relatively consistent lifespan extension under caloric restriction, despite the broad phylogenetic range represented by the species included in our study.

**Figure 3.**
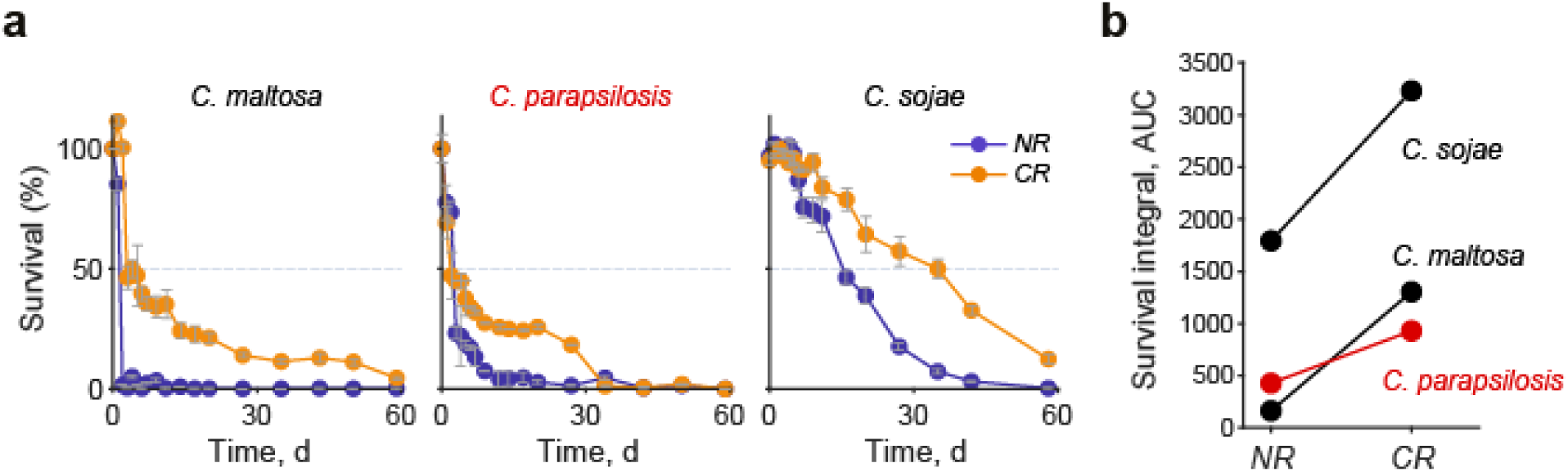
CFU-based measurements recapitulate interspecific CLS differences and responses to caloric restriction. **(a)** CFU-based CLS profiles for *C. maltosa*, *C. parapsilosis*, and *C. sojae* under non-restricted (NR, 2% glucose) and caloric-restriction conditions (CR, 0.5% glucose). Points represent mean survival across three independent experiments, and error bars indicate the SEM at each time point. **(b)** Reaction norms showing the change in survival integral from NR to CR for each species.

We next asked whether this CLS variation was related to variation in pathogenicity across species. Clinical status was defined according to the WHO fungal priority pathogens list, distinguishing listed from non-listed species (**Table 1**). For comparative purposes, WHO-listed species were treated as the pathogenic group. WHO-listed *Candida* were interspersed with non-listed species across the phylogeny, providing no evident phylogenetic grouping by WHO priority status (**Fig. 2b**) (Gabaldón et al., 2016). CLS profiles did not segregate clearly by this classification, although WHO-listed species occupied intermediate and high ranges of survival integrals under NR and exhibited similarly strong lifespan extension under CR (**Fig. 2c**). The non-lisited species, however, spanned a much broader range of CLS phenotypes and responses. *C. sojae* was already long-lived under NR and further extended survival under CR, whereas *C. maltosa* had a much shorter baseline CLS under NR but showed a comparably strong response to CR. *C. dubliniensis* differed from both, combining short baseline survival with the weakest CR response among the *Candida* species examined.

Notably, although *C. dubliniensis* is not included in the WHO priority list, it is a common inhabitant of the human body, whereas *C. sojae* and *C. maltosa* are primarily associated with industrial settings. Thus, although WHO-listed species occupied a more restricted range of CLS profiles, the long-lived phenotype was not specific to WHO-listed species.

Together, these results show that CLS varies extensively across *Candida* species, whereas extension of survival by CR was observed in every species examined and was generally consistent in magnitude despite large differences in baseline NR survival. WHO-listed species tended to occupy the intermediate to long-lived portion of the CLS distribution, but extended CLS was not restricted to clinically prioritized species, as illustrated by the long-lived *C. sojae*.

### 3.3. Longer chronological lifespan is associated with greater virulence within ***Candida albicans***

Given that pathogenic species tended toward longer CLS but long lifespan was not restricted to them, we next asked whether quantitative variation in stationary-phase survival was more closely associated with virulence within a single species. To this end, we analyzed a panel of 20 genetically diverse *C. albicans* strains whose virulence had been previously characterized in a murine model of systemic infection (Wu et al., 2007). Virulence estimates were derived from the published host survival data and expressed as a virulence score calculated as 1 minus the normalized area under the host survival curve, such that higher scores corresponded to greater virulence. CLS was measured by flow cytometry at intermediate aging time points that captured substantial inter-strain variation in survival (days 10 and 15; **Fig. 4a,b**).

**Figure 4.**
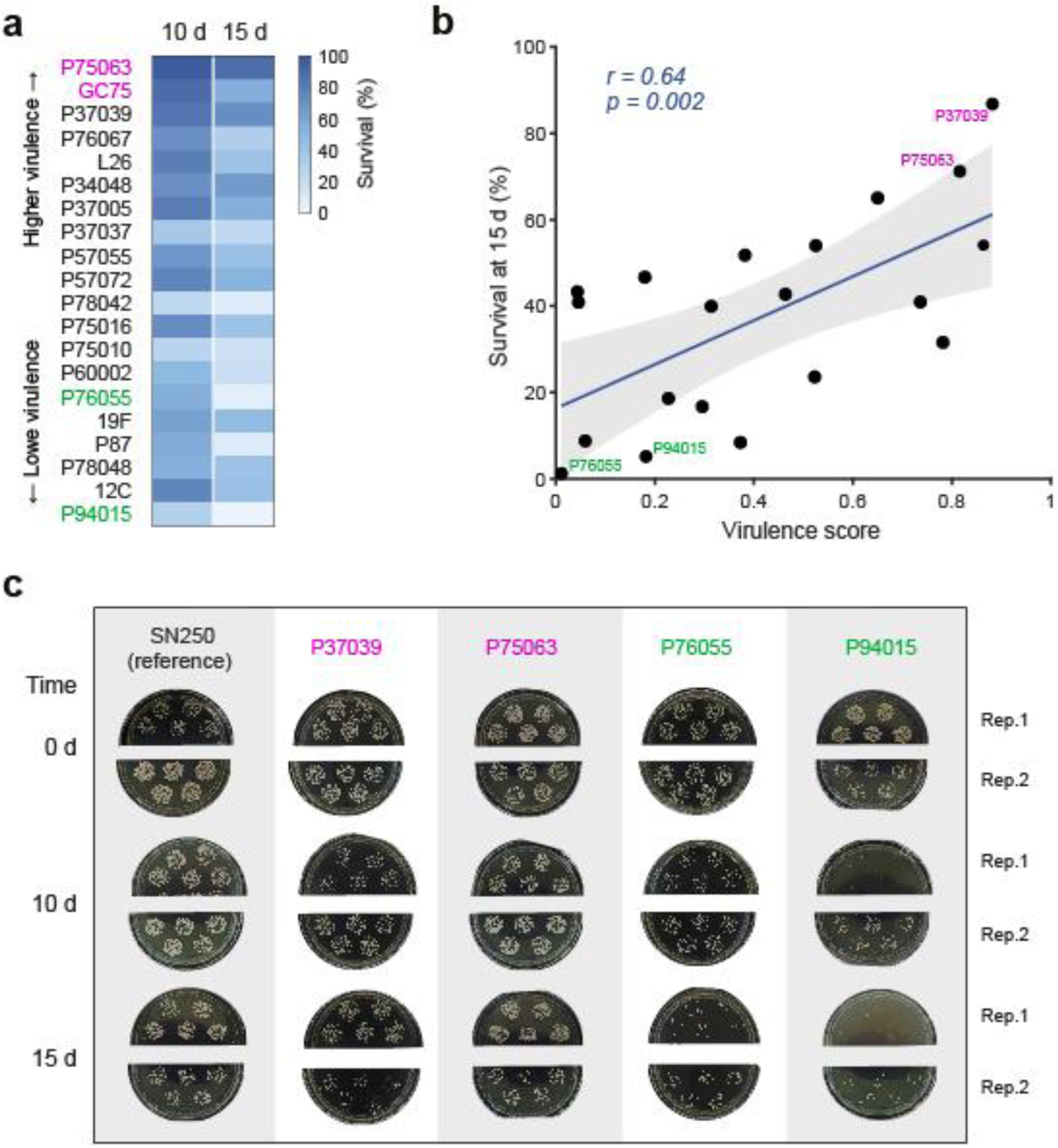
Chronological lifespan is associated with virulence among *Candida albicans* strains. **(a)** Flow cytometry-based survival of 20 *C. albicans* strains after 10 and 15 days of chronological aging. Strains are ordered according to virulence estimates derived from previously published murine systemic-infection data (Wu et al., 2007), using a virulence score calculated as 1 minus the normalized area under the host survival curve; higher scores correspond to greater virulence. **(b)** Correlation between CLS and virulence across the strain panel at day 15. Each point represents one strain; the solid line indicates the linear regression fit and the shaded area the 95% CI. Pearson’s correlation coefficient and corresponding *p*-value are shown. **(c)** Representative CFU plates from the two highest-ranked (pink labels) and two lowest-ranked (green labels) strains along with the reference strain, based on flow cytometry-derived CLS. Two replicate plates at days 0, 10, and 15 of chronological aging are shown.

CLS varied at both 10 and 15 days of stationary-phase from near-complete loss of viability to high survival (**Fig. 4a**). This variation was associated with virulence, with strains maintaining greater survival during chronological aging tending to be more virulent, as reflected by higher virulence scores (**Fig. 4b**). Statistically, CLS survival was positively correlated with virulence score at both day 10 (*r* = 0.59, *p* = 0.006) and day 15 (*r* = 0.64, *p* = 0.002; Pearson correlations). These results show that stationary-phase survival is positively associated with virulence among *C. albicans* strains.

To determine whether the strain differences identified by flow cytometry were also reflected in colony-forming capacity, we selected the two longest-lived and two shortest-lived strains based on their flow cytometry profiles and measured their CLS independently by CFU (**Fig. 4c**). Strains classified as long-lived by flow cytometry retained greater colony-forming capacity during aging than strains classified as short-lived, supporting the survival differences detected by the membrane integrity-based assay. Together, these results identify an association between extended stationary-phase survival and greater virulence within *C. albicans*, whereas across species long CLS was not restricted to WHO-listed pathogenic species.

## 4. Discussion

Chronological lifespan has provided a powerful model for understanding survival of non-dividing cells in *S. cerevisiae*, where studies have linked this trait with nutrient availability, stress resistance, and cellular maintenance (Cruz-Bonilla et al., 2025; Fabrizio and Longo, 2007; Smith et al., 2016). However, its relevance across human opportunistic yeast pathogens remains poorly understood. Our comparative analysis across *Candida* species revealed extensive baseline NR variation in stationary-phase survival together with a shared response to reduced glucose: caloric restriction extended CLS in every species examined, despite the large evolutionary distances among them. Pathogenic species tended toward longer CLS, but long lifespan was not restricted to them, whereas within *C. albicans* longer CLS correlated with greater virulence. Parallel CFU and flow-cytometry measurements further showed that colony-forming capacity and membrane integrity provide overlapping but non-equivalent readouts of survival, particularly under caloric restriction. Together, these findings position CLS as a highly variable but broadly nutrient-responsive trait whose relationship with pathogenicity differs between interspecific and intraspecific comparisons.

The divergence between CFU- and membrane integrity-based measurements under caloric restriction emphasizes that chronological survival is not a single physiological state. Classical CFU-based CLS assays define survival through the ability of aged cells to resume proliferation, whereas PI exclusion reports maintenance of membrane integrity without requiring subsequent growth (Tang et al., 2025). Our results indicate that under standard conditions these properties decline together, but under CR nutrient limitation can generate cells whose membrane remains intact while no longer contributing to the colony-forming population. Studies using fluorescence-based viability measurements in stressed yeasts have similarly identified heterogeneous or intermediate physiological states that are not necessarily equivalent to proliferative competence (Kwolek-Mirek and Zadrag-Tecza, 2014; Tang et al., 2025). Thus, the two assays should be viewed as complementary rather than interchangeable readouts of CLS. This distinction becomes particularly important in comparative studies, where environmental conditions may differentially affect proliferative competence and other components of cellular viability.

Our assays focused on measuring CLS of yeast-form populations as a first approach to understanding stationary-phase survival in yeasts of the CTG clade. Although both yeast and filamentous morphologies are biologically important for these species, particularly during host colonization and infection (Kadosh and Mundodi, 2020; Witchley et al., 2019), we focused initially on the yeast form because a consistent morphological state facilitates reproducible survival measurements. Filamentation introduces variation in cell size, shape, aggregation, and cellular organization, which can compromise both CFU enumeration and flow-cytometric assessment of membrane integrity. Shifts in the relative abundance of yeast and filamentous cells during aging could therefore confound CLS measurements. Previous work using replicative lifespan reported similar longevity in yeast and filamentous forms of *C. albicans* (Fu et al., 2008). Although replicative lifespan measures mitotic survival rather than chronological survival during stationary phase, this observation raises the possibility that lifespan may not differ strongly between morphologies. Future studies that overcome the challenges of following filamentous cells over time, potentially using microfluidic approaches, will help determine whether this also applies to CLS.

The variation in CLS across *Candida* species suggests that stationary-phase survival is a highly variable trait, with substantial differences even among related species. Phenotypic surveys of *Candida* have likewise revealed extensive interspecies and intraspecific variation in stress resistance, growth, and metabolic traits (Beyer et al., 2026; Brandt et al., 2023). In contrast to this baseline NR diversity, caloric restriction extended CLS in every species examined, with broadly similar responses across most of the panel. Baseline longevity and responsiveness to nutrient limitation therefore appear to represent partially separable components of stationary-phase physiology. *C. sojae* combines relatively long baseline CLS with a strong response to CR, whereas *C. maltosa* is comparatively short-lived but responds similarly strongly, and *C. dubliniensis* combines short baseline survival with a weaker CR response. Importantly, the broad phenotypic range among non-pathogenic species, including the long CLS of *C. sojae*, shows that prolonged stationary-phase survival can occur in species not associated with human infection and is therefore not a pathogen-specific trait. The consistency of the CR response across species is particularly notable because responses to CR can vary substantially among genotypes within a species. Natural *S. cerevisiae* populations show genotype- and diet-dependent variation in CLS (Jung et al., 2018), while effects of CR on replicative lifespan range from extension to little effect or even lifespan shortening across genetic backgrounds (McLean et al., 2024; Schleit et al., 2013). Thus, the broadly shared response observed across our *Candida* species contrasts with substantial intraspecific heterogeneity reported in other systems. Whether variability in CR responsiveness differs systematically across evolutionary scales will require broader sampling.

The contrasting relationships between CLS and pathogenicity across and within species are particularly informative. Across *Candida* species, pathogenic and non-pathogenic species occupy overlapping portions of the CLS distribution. This is consistent with the view that fungal pathogenicity is a complex phenotype arising from multiple traits affecting growth, stress tolerance, host interaction, and responses to the environment (Priest and Lorenz, 2015; Rokas, 2022). Within *C. albicans*, however, strains maintaining higher survival during chronological aging were also more virulent. *C. albicans* populations are themselves genetically and phenotypically diverse, with substantial strain-to-strain variation in traits including stress responses and virulence (Hirakawa et al., 2015; Shankar et al., 2020). These patterns suggest that the relationship depends on evolutionary scale, with CLS associated with virulence within *C. albicans* but not with pathogenicity across more deeply diverged *Candida* species. Across deeper evolutionary divergence, CLS is embedded within many other species-specific traits that contribute to host interaction and pathogenic potential; within a single species, where much of this biological background is shared, quantitative variation in stationary-phase survival may become more closely coupled to variation in virulence.

The intraspecific association between CLS and virulence raises the question of which cellular and physiological processes contribute to both traits. Entry into stationary phase in yeast involves extensive metabolic remodeling, induction of stress-response programs, and maintenance processes that promote survival during prolonged nutrient limitation (Breeden and Tsukiyama, 2022; den Ridder et al., 2023; Sun and Gresham, 2021; Werner-Washburne et al., 1993). In *Candida* species, non-proliferative or slowly proliferating states have also been associated with increased tolerance to oxidative, thermal, and nutritional stresses (Cuéllar-Cruz et al., 2008; Kusch et al., 2008; Richards et al., 2010). Natural variation in CLS has been linked to evolutionary history in *S. cerevisiae*, where domestication was modestly associated with longer CLS but lifespan variation remained predominantly clade-specific (De Chiara et al., 2022). This raises the possibility that stationary-phase survival can become associated with adaptation to selective environments without being specific to any one lifestyle. Such properties could be relevant during infection, where fungal cells encounter nutrient limitation, immune-mediated stress, physicochemical challenges, and antifungal exposure (Alves et al., 2020; Pountain et al., 2021). Variation in these underlying processes could therefore generate the observed covariance between CLS and virulence without chronological aging itself being causal. Aging-associated physiological traits have previously been implicated in fungal persistence and stress tolerance (Bhattacharya et al., 2019), although a direct intraspecific relationship between CLS and virulence in *Candida* has, to our knowledge, not been established. Moreover, CLS was measured in a defined laboratory environment, whereas virulence was derived from a systemic murine infection model; these contexts differ substantially. Identifying whether particular stress-response, metabolic, or maintenance pathways contribute to both phenotypes will require targeted perturbations to determine whether the correlation reflects shared mechanisms rather than merely correlated strain variation.

## 5. Conclusion

This study reveals two contrasting features of CLS variation in *Candida*. Across species, baseline lifespan under non-restricted (NR) conditions is highly diverse, yet caloric restriction produces a shared extension of survival, indicating that responsiveness to reduced glucose is retained despite substantial differences in stationary-phase longevity. Long lifespan itself is not restricted to pathogenic species. Within *C. albicans*, however, longer CLS correlates with greater virulence, pointing to physiological determinants that may contribute to both traits without establishing a causal role for chronological aging in infection. Defining those shared determinants through targeted genetic and physiological perturbations will be necessary to understand why CLS and virulence covary among *C. albicans* strains.

## Acknowledgements

We are grateful to Jimena Meneses-Plascencia, Susana Ruiz-Castro, and Mayra I. Flores Barraza for skillful technical assistance and to Diana Ascencio for critical reading of the manuscript. This work was funded by Secretaría de Ciencia, Humanidades, Tecnología e Innovación (Secihti) grants CF-2023-G-695 and CBF-2026-3450.

## CRediT authorship contribution statement

**Alberto Luna-Carrera**: Conceptualization, Methodology, Investigation, Formal analysis, Visualization, Writing – original draft. **Lilian Rodríguez-Flores**: Investigation. **Ericka Moreno-Méndez**: Methodology, Investigation. **Eugenio Mancera**: Conceptualization, Funding acquisition, Supervision, Resources, Writing – review & editing. **Alexander DeLuna**: Conceptualization, Formal analysis, Funding acquisition, Supervision, Resources, Writing – original draft. All authors read and critically evaluated the final version of the manuscript.

## AI-assisted technologies in manuscript preparation

During the preparation of the revised version of this manuscript the authors used ChatGPT (*GPT-5.5*, OpenAI) for language editing and proofreading assistance. After using this tool, the authors reviewed and edited all content as needed and take full responsibility for the content of the published article.

